## Supplemental Information for "Dominance in dogs as rated by owners corresponds to ethologically valid markers of dominance"

**How do owners perceive dominance in dogs?**

### Correlations of the items

**Methods**

We used the binary behavioural categories: which of your dogs’ expresses the behaviour more, the "dominant" = 1 or the "subordinate" = 0, of different items: bark, lick mouth, eat first, reward, fight, play ball, greet owner, walk first, resting place, overmark, defend group, smarter, obedient, aggressive, impulsive, size, physical condition and age, and correlated them using a Pearson Correlation. For this analysis we only used dyads where we had no missing information (N=215).

**Results and Discussion**

The results of the correlation are displayed in Table S1. The analysis revealed many small correlations (47 at 0.1 to 0.3), four medium correlations (0.3 to 0.5) and one large association (0.5 to 1.0). Here we shall focus mainly on nine items (as these were the most related to dominance status): Which dog starts to bark first, licks the mouth of the other, eats first, obtains the reward, wins fights, walks at the front, acquires the better resting place, defends the group, and is more aggressive. Dogs that bark first were also more likely to defend the group and were more aggressive. Dogs that licked the mouth of the other won fights less often. This negative correlation was expected as licking the mouth of the other individual occurs more often in subordinate individuals. Dogs that eat first were more often rewarded first (r>0.5), and dogs that were rewarded first more often won fights and obtained the best resting place. Dogs that won fights obtained the better resting place and defended the group. Individuals that walked at the front or first obtained the better resting place and defended the group. Finally, dogs that defended the group were more aggressive.

When we consider only the more significant associations among the other items, unsurprisingly, individuals who played with the ball were perceived by the owner as being in a better physical condition, more aggressive individuals were more impulsive, and older individuals were less impulsive and in worse physical condition. The large association found was between eat first and reward, which was expected, as dogs that were more motivated to obtain food are more likely to receive a reward from the owner before an individual who is not as food motivated.

| **Item** | **Bark** | **Lick mouth** | **Eat first** | **Reward** | **Fight** | **Play ball** | **Greet owner** | **Walk first** | **Resting place** | **Overmark** | **Defend group** | **Smarter** | **Obedient** | **Aggressive** | **Impulsive** | **Size** | **Physical condition** | **Age** |
| --- | --- | --- | --- | --- | --- | --- | --- | --- | --- | --- | --- | --- | --- | --- | --- | --- | --- | --- |
| **Bark** | 1 | .015 | .007 | -.088 | -.071 | .085 | .086 | .046 | -.112 | .085 | .134^*^ | -.079 | -.125 | .260^**^ | .152^*^ | -.097 | -.048 | .077 |
| **Lick mouth** | .015 | 1 | -.001 | -.062 | -.183^**^ | .110 | .084 | -.063 | -.064 | -.152^*^ | -.088 | -.073 | -.011 | -.047 | .192^**^ | -.106 | .082 | -.249^**^ |
| **Eat first** | .007 | -.001 | 1 | .556^**^ | .081 | .152^*^ | .040 | .002 | .104 | .022 | -.044 | -.091 | -.131 | -.024 | .006 | .227^**^ | .221^**^ | .006 |
| **Reward** | -.088 | -.062 | .556^**^ | 1 | .217^**^ | .037 | .075 | .115 | .184^**^ | .105 | .019 | -.017 | -.180^**^ | -.020 | -.046 | .261^**^ | .177^**^ | .039 |
| **Fight** | -.071 | -.183^**^ | .081 | .217^**^ | 1 | .035 | .011 | .018 | .197^**^ | .021 | .235^**^ | .151^*^ | .110 | -.012 | -.011 | .210^**^ | .128 | .023 |
| **Play ball** | .085 | .110 | .152^*^ | .037 | .035 | 1 | .093 | -.139^*^ | -.079 | -.014 | .161^*^ | .134 | .108 | -.049 | .121 | .022 | .310^**^ | -.158^*^ |
| **Greet owner** | .086 | .084 | .040 | .075 | .011 | .093 | 1 | .067 | .054 | .089 | .076 | .134^*^ | .125 | -.138^*^ | .184^**^ | -.021 | .182^**^ | -.160^*^ |
| **Walk first** | .046 | -.063 | .002 | .115 | .018 | -.139^*^ | .067 | 1 | .229^**^ | .231^**^ | .146^*^ | .095 | -.103 | .032 | .094 | .025 | .147^*^ | -.064 |
| **Resting place** | -.112 | -.064 | .104 | .184^**^ | .197^**^ | -.079 | .054 | .229^**^ | 1 | .165^*^ | .036 | .120 | -.059 | -.037 | .139^*^ | -.038 | .054 | -.010 |
| **Overmark** | .085 | -.152^*^ | .022 | .105 | .021 | -.014 | .089 | .231^**^ | .165^*^ | 1 | .237^**^ | .013 | -.123 | .047 | .039 | .125 | .092 | -.032 |
| **Defend group** | .134^*^ | -.088 | -.044 | .019 | .235^**^ | .161^*^ | .076 | .146^*^ | .036 | .237^**^ | 1 | .162^*^ | .009 | .187^**^ | .061 | .094 | .094 | .045 |
| **Smarter** | -.079 | -.073 | -.091 | -.017 | .151^*^ | .134 | .134^*^ | .095 | .120 | .013 | .162^*^ | 1 | .292^**^ | -.146^*^ | -.131 | .108 | .097 | .145^*^ |
| **obedient** | -.125 | -.011 | -.131 | -.180^**^ | .110 | .108 | .125 | -.103 | -.059 | -.123 | .009 | .292^**^ | 1 | -.268^**^ | -.277^**^ | .220^**^ | .003 | .118 |
| **Aggressive** | .260^**^ | -.047 | -.024 | -.020 | -.012 | -.049 | -.138^*^ | .032 | -.037 | .047 | .187^**^ | -.146^*^ | -.268^**^ | 1 | .347^**^ | -.215^**^ | -.039 | -.071 |
| **Impulsive** | .152^*^ | .192^**^ | .006 | -.046 | -.011 | .121 | .184^**^ | .094 | .139^*^ | .039 | .061 | -.131 | -.277^**^ | .347^**^ | 1 | -.222^**^ | .247^**^ | -.389^**^ |
| **Size** | -.097 | -.106 | .227^**^ | .261^**^ | .210^**^ | .022 | -.021 | .025 | -.038 | .125 | .094 | .108 | .220^**^ | -.215^**^ | -.222^**^ | 1 | .269^**^ | .085 |
| **Physical condition** | -.048 | .082 | .221^**^ | .177^**^ | .128 | .310^**^ | .182^**^ | .147^*^ | .054 | .092 | .094 | .097 | .003 | -.039 | .247^**^ | .269^**^ | 1 | -.317^**^ |
| **Age** | .077 | -.249^**^ | .006 | .039 | .023 | -.158^*^ | -.160^*^ | -.064 | -.010 | -.032 | .045 | .145^*^ | .118 | -.071 | -.389^**^ | .085 | -.317^**^ | 1 |

**Table S1.** Results of the Pearson correlation of the 20 items showing the direction and magnitude of effects and the significance level of the terms.

* Correlation is significant at the 0.05 level (2-tailed). ** Correlation is significant at the 0.01 level (2-tailed). Sample size = 215 dyads.

### Differences in the number of dominance related behaviours expressed in dominants and subordinates

**Methods**

*Calculating the Dominance Score and the Difference Score*

We created a “dominance score” by summing all the items that were significantly associated with a “dominant” status (bark, lick mouth, eat first, reward, fight, walk first, resting place, pee, defend group, smart, aggressive, and impulsive) for each dog in every dyad. Then we created a “difference score” by subtracting the "subordinates" dominance score from the “dominants” for each dyad. Because we obtained some negative values, and the statistical model work only with positive values, we added a constant (12) to the difference score. After a power transformation to achieve normal distribution (Boxcox, lamda = 0.67, three outliers removed) the difference score was then used as the response variable in a General linear model that was performed in R, to identify the key variables associated with the difference score. All possible interactions between dominant sex (male or female), subordinate sex (male or female), dominant neuter status (intact or neutered), subordinate neuter status (intact or neutered) and dominant age (older or younger) were entered into the model. We also included the main effect of the order the dogs were entered into the questionnaire (Dog A first or second) to examine order effects. We included only the dyads where an asymmetry in dominance was detected by the owner (N= 931).

**Results and Discussion**

“Dominants” had higher dominance difference score (“dominant” mean ± SD = 6.03±2.46, range = 0-12 vs. 2.92±1.91, range = 0 – 8, Mann-Whitney U test = -27.326, P <0.001). In comparison, dogs that were rated as similar in status had similar scores (3.23 ± 2.01, range = 0 – 9 for Dog A, and 3.46 ± 2.10, range = 0 – 11 for Dog B, Mann-Whitney U test = 0.837, P =0.403). However, there was quite some variation in both “dominant” and “subordinate” individuals, which reflects the complex nature of social relationships, and the influence of context and previous experience on behaviour.

|  | Estimate | Standard error | T value | F | P | Partial eta |
| --- | --- | --- | --- | --- | --- | --- |
| Order: Dog B | -0.407 | 0.138 | -2.941 | 8.4453 | 0.004* | 0.009 |
| Dom. sex: Male | 0.118 | 0.136 | 0.866 | 0.7491 | 0.3870 | 0.001 |
| Sub. sex: Male | -0.613 | 0.161 | -3.806 | 14.487 | <0.001* | 0.015 |
| Sub. Neut. status: Neutered | -0.305 | 0.135 | -2.262 | 5.119 | 0.024* | 0.006 |
| Dom age: Younger | 0.324 | 0.126 | 2.568 | 6.5948 | 0.010* | 0.007 |
| Dom. Sex:Sub sex-Male:Male | 0.413 | 0.195 | 2.116 | 4.476 | 0.035* | 0.005 |
| Sub. Sex:Sub. Neut. Status- Male:Neutered | 0.434 | 0.195 | 2.221 | 4.934 | 0.027* | 0.005 |

**Table S2.** Results of the general linear model showing the direction and magnitude of effects and the significance level of the terms in the demographic variables associated with “Difference score”. Significant P values (indicated with *) indicate which group differs from the reference value in the respective analysis.

Results of the general linear model utilising the difference score as the response variable revealed significant main effects of subordinate sex, subordinate neuter status, order, and dominant age (Table S2). However, the overall variance explained by the model was low (Multiple R-squared: 0.03601, F-statistic: 4.925 on 7 and 923 DF, P < 0.001). Two of the interactions were significant after model reduction, dominant sex:subordinate sex, and subordinate sex:subordinate neuter status. The mean difference score when the dominant sex was female and the subordinate was male was lower than all other sex combinations (female female, male female and male male; Figure S1), regardless of neuter status. In addition, when the subordinate was an intact female, the difference score was higher than all other subordinate sex neuter status combinations (female neutered, male intact and male neutered; Figure S2).

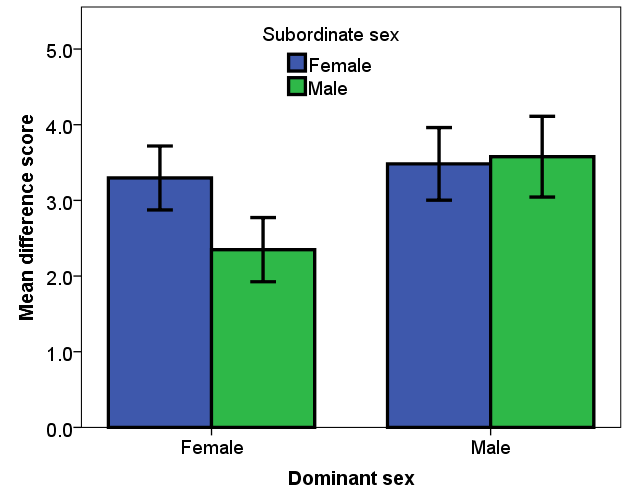

**Figure S1.** Mean and 95% confidence intervals of the difference score of the four dominant sex and subordinate sex combinations (female female, female male, male female and male male). The interaction was significant at P < 0.05.

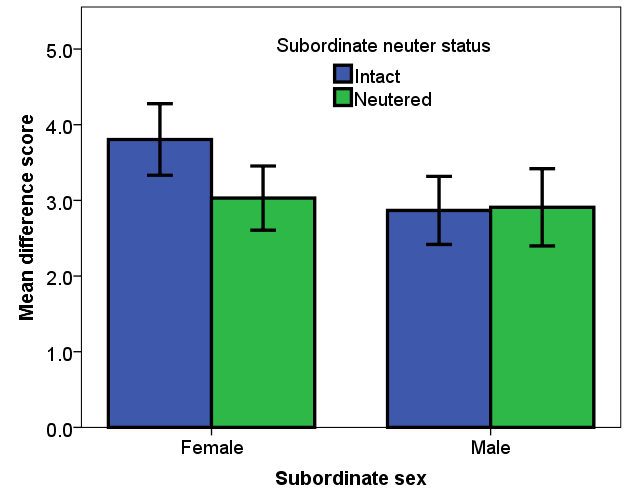

**Figure S2.** Mean and 95% confidence intervals of the difference score of the four subordinate sex and subordinate neuter status combinations (female intact, female neutered, male intact and male neutered). The interaction was significant at P < 0.05.

In general, the mean difference score was higher when the subordinate was female compared to male (Figure S3), and when the subordinate was intact compared to neutered (Figure S4). Our results partially collaborate the suggestion that mixed sex dog dyads tend to have more defined relationships, in comparison to same sex pairs (McGreevy et al., 2012). However, when the female of the pair was viewed as dominant, the difference score was much lower, as male dominants received a higher dominance score than female dominants in mixed sex groups. Which points to a possible sex influence on some dominance related behaviours. In the current study, when a male was dominant in a mixed sex pair, he more often marked over his female partner and defended the group in case of perceived danger, than when a female was dominant. Dominant males may be performing mate-guarding behaviour in an attempt to control intact females mating opportunities. This behaviour includes increased urine marks on or near a females urine spots, which serves to hide the odour trail of an oestrous female from other dogs (Dunbar & Buehler, 1980). When female free-ranging dogs are in heat, males tend to become more aggressive towards each other, and hierarchies become more pronounced during this time (Daniels, 1983). However, this does not explain why for some owners of intact mixed sex dyads the female was perceived as dominant. There is evidence from humans that socially dominant males and females are more similar in behavioural profiles (regardless of age), than is commonly believed (Hawley, Little & Card, 2008). Biologists have underrated overt competitiveness in females, as evolutionary and biological approaches suggested that social dominance is predominately an aspect of male social organization. However, it is also possible that the owners reinforced the position of the female over the male (allowing only the female to get the best sleeping position, treats, and rewards etc.), either as they preferred that individual, or they were seen as weaker and therefore needed protection from the larger male.

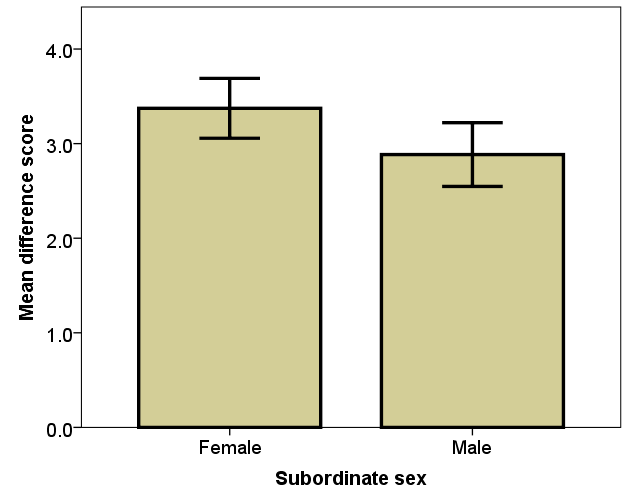

**Figure S3.** Mean and 95% confidence intervals of the difference score of female subordinates in comparison to male subordinates (significant at P < 0.05).

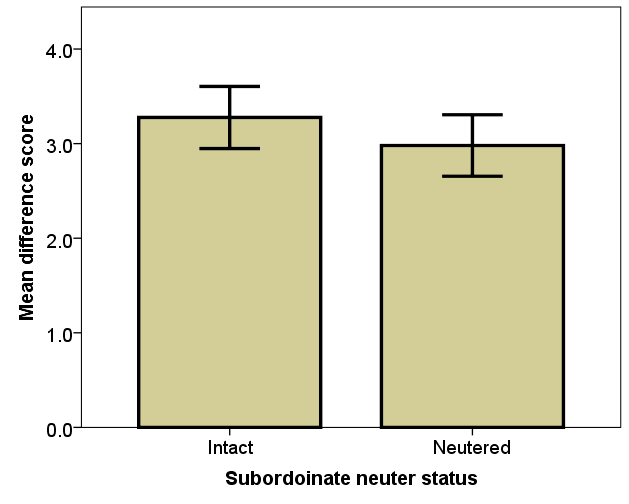

**Figure S4.** Mean and 95% confidence intervals of the difference score of intact subordinates in comparison to neutered subordinates (significant at P < 0.05).

The results of the model indicate that it is actually the subordinate identity rather than the dominant that is more important when examining the difference score. Regardless of the dominant identity, when the subordinate was female they showed fewer dominance related behaviours, especially if they were reproductively intact. Lower ranking females and intact individuals (regardless of sex) in general may show more submissive behaviours in order to remain within the group without conflict. Previous studies have determined that subordinates play a central role in establishing dominance relationships as the outcome of dyadic interactions are often determined by the subordinate dogs’ behaviour (Cafazzo et al., 2010).

Interestingly, when the dominant was the younger animal the difference score was higher, regardless of dyad combination (Table S1, Figure S5). In the current study, younger individuals were described as more impulsive and in better physical condition than older individuals. In addition, due to higher activity levels than the older individual they may appear to take on some of the behaviours of dominance, such as walking first and defending the group, as well as being quicker to eat first and obtain rewards. Interestingly, when we examined the differences between older and younger dominant individuals, we found that when the dominant was younger than the subordinate, they licked the mouth of the subordinate more often than when the dominant was the older animal (younger dominant proportion lick mouth = 0.64, older dominant = 0.22, proportion difference = 0.42, Z = 10.72, P<0.001). Mouth licking is a sign of formal dominance. Licking the mouth of the partner, usually during greeting ceremonies, signals their acceptance of lower social status, (Bonanni et al., 2010), which puts the real identity of the dominant individual into question.

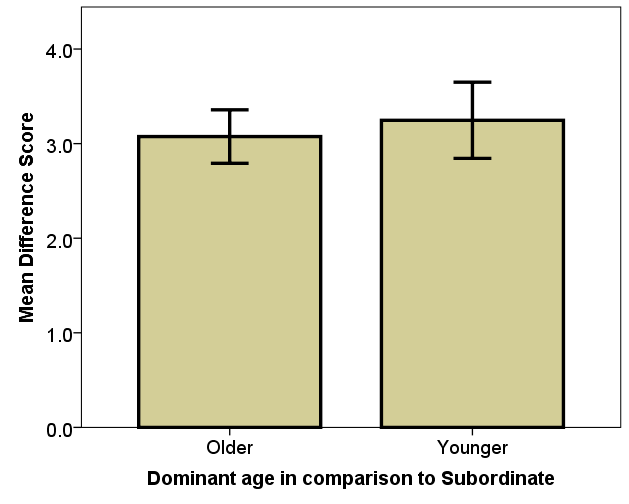

**Figure S5.** Mean and 95% confidence intervals of the difference score of older and younger dominants (in comparison to the subordinate age) (significant at P < 0.05).

There was also a significant order effect. “Dominant” Dog A-s differed more from their partners than “Dominant” Dog B-s (Table S6). This indicates that when owners were more confident of the differences between the two dogs in their household, they tended to put the more dominant individual first (i.e. as “Dog A”).

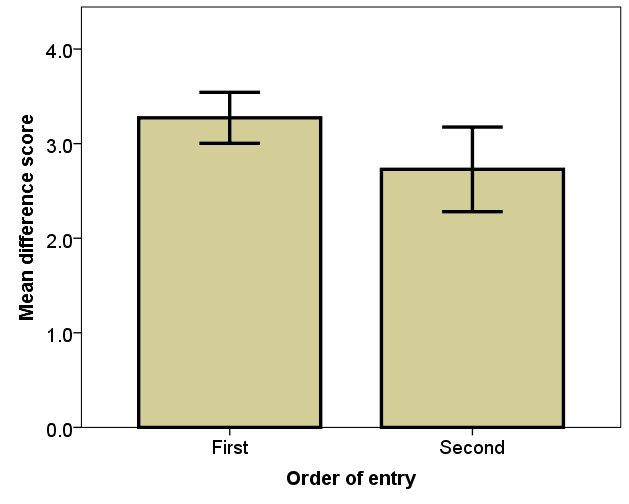

**Figure S6.** Mean and 95% confidence intervals of the difference score of the order of entry (the dominant dog was entered into the questionnaire first, compared to second) (significant at P < 0.05).

### Difference score models R code

**Full Model**

```
> B <- lm(bcPower(diff,0.67) ~ order+domsex*subsex*domneut*subneut*domage, data=newdomdiff, na.action = na.omit)
> summary(B)

Call:
lm(formula = bcPower(diff, 0.67) ~ order + domsex * subsex * domneut *
    subneut * domage, data = newdomdiff, na.action = na.omit)

Residuals:
    Min      1Q  Median      3Q     Max
-4.1361 -1.0058  0.0144  0.9924  3.8629

Coefficients:
                                                                    Estimate Std. Error t value Pr(>|t|)
(Intercept)                                                          7.93873    0.19459  40.796  <2e-16 ***
orderSecond                                                         -0.40616    0.14179  -2.865  0.00427 **
domsexmale                                                           0.13395    0.32269   0.415  0.67817
subsexmale                                                          -0.85396    0.32936  -2.593  0.00967 **
domneutneutered                                                     -0.16682    0.32588  -0.512  0.60884
subneutneutered                                                      0.03864    0.55035   0.070  0.94404
domageyounger                                                        0.12069    0.32851   0.367  0.71343
domsexmale:subsexmale                                                0.59479    0.45375   1.311  0.19024
domsexmale:domneutneutered                                          -0.04079    0.49548  -0.082  0.93441
subsexmale:domneutneutered                                           0.24005    0.46128   0.520  0.60292
domsexmale:subneutneutered                                          -0.28115    0.65990  -0.426  0.67018
subsexmale:subneutneutered                                           0.74239    0.72442   1.025  0.30573
domneutneutered:subneutneutered                                     -0.37325    0.62916  -0.593  0.55316
domsexmale:domageyounger                                             0.44765    0.57563   0.778  0.43697
subsexmale:domageyounger                                             0.77942    0.56961   1.368  0.17155
domneutneutered:domageyounger                                        0.64607    0.64043   1.009  0.31334
subneutneutered:domageyounger                                       -0.19871    0.73811  -0.269  0.78783
domsexmale:subsexmale:domneutneutered                                0.59403    0.67749   0.877  0.38082
domsexmale:subsexmale:subneutneutered                               -0.45324    0.93002  -0.487  0.62614
domsexmale:domneutneutered:subneutneutered                           0.28787    0.80600   0.357  0.72106
subsexmale:domneutneutered:subneutneutered                          -0.03642    0.83171  -0.044  0.96508
domsexmale:subsexmale:domageyounger                                 -1.54094    0.80672  -1.910  0.05644 .
domsexmale:domneutneutered:domageyounger                            -0.85590    1.00677  -0.850  0.39547
subsexmale:domneutneutered:domageyounger                            -1.10279    0.86239  -1.279  0.20131
domsexmale:subneutneutered:domageyounger                            -0.29191    0.97812  -0.298  0.76544
subsexmale:subneutneutered:domageyounger                            -1.17636    1.02687  -1.146  0.25227
domneutneutered:subneutneutered:domageyounger                        0.15012    0.96147   0.156  0.87596
domsexmale:subsexmale:domneutneutered:subneutneutered               -0.71207    1.14007  -0.625  0.53240
domsexmale:subsexmale:domneutneutered:domageyounger                  1.44368    1.30015   1.110  0.26712
domsexmale:subsexmale:subneutneutered:domageyounger                  3.04748    1.67442   1.820  0.06909 .
domsexmale:domneutneutered:subneutneutered:domageyounger             0.57975    1.36469   0.425  0.67107
subsexmale:domneutneutered:subneutneutered:domageyounger             0.91631    1.27713   0.717  0.47327
domsexmale:subsexmale:domneutneut:subneutneut:domageyoung	       -2.28249    2.08789  -1.093  0.27460
---
Signif. codes:  0 ‘***’ 0.001 ‘**’ 0.01 ‘*’ 0.05 ‘.’ 0.1 ‘ ’ 1

Residual standard error: 1.456 on 898 degrees of freedom
Multiple R-squared:  0.05884,	Adjusted R-squared:  0.0253
F-statistic: 1.754 on 32 and 898 DF,  p-value: 0.006364


> etasq(B, anova = TRUE, type=3)
Anova Table (Type III tests)

Response: bcPower(diff, 0.67)
                                     Partial eta^2 Sum Sq  Df   F value    Pr(>F)
(Intercept)                                0.64954 3526.9   1 1664.3441 < 2.2e-16 ***
order                                      0.00906   17.4   1    8.2057  0.004273 **
domsex                                     0.00019    0.4   1    0.1723  0.678166
subsex                                     0.00743   14.2   1    6.7226  0.009675 **
domneut                                    0.00029    0.6   1    0.2620  0.608845
subneut                                    0.00001    0.0   1    0.0049  0.944041
domage                                     0.00015    0.3   1    0.1350  0.713426
domsex:subsex                              0.00191    3.6   1    1.7183  0.190245
domsex:domneut                             0.00001    0.0   1    0.0068  0.934408
subsex:domneut                             0.00030    0.6   1    0.2708  0.602922
domsex:subneut                             0.00020    0.4   1    0.1815  0.670176
subsex:subneut                             0.00117    2.2   1    1.0502  0.305732
domneut:subneut                            0.00039    0.7   1    0.3520  0.553160
domsex:domage                              0.00067    1.3   1    0.6048  0.436970
subsex:domage                              0.00208    4.0   1    1.8723  0.171549
domneut:domage                             0.00113    2.2   1    1.0177  0.313338
subneut:domage                             0.00008    0.2   1    0.0725  0.787829
domsex:subsex:domneut                      0.00086    1.6   1    0.7688  0.380824
domsex:subsex:subneut                      0.00026    0.5   1    0.2375  0.626137
domsex:domneut:subneut                     0.00014    0.3   1    0.1276  0.721059
subsex:domneut:subneut                     0.00000    0.0   1    0.0019  0.965080
domsex:subsex:domage                       0.00405    7.7   1    3.6486  0.056435 .
domsex:domneut:domage                      0.00080    1.5   1    0.7228  0.395468
subsex:domneut:domage                      0.00182    3.5   1    1.6352  0.201311
domsex:subneut:domage                      0.00010    0.2   1    0.0891  0.765438
subsex:subneut:domage                      0.00146    2.8   1    1.3124  0.252274
domneut:subneut:domage                     0.00003    0.1   1    0.0244  0.875961
domsex:subsex:domneut:subneut              0.00043    0.8   1    0.3901  0.532401
domsex:subsex:domneut:domage               0.00137    2.6   1    1.2330  0.267124
domsex:subsex:subneut:domage               0.00368    7.0   1    3.3125  0.069089 .
domsex:domneut:subneut:domage              0.00020    0.4   1    0.1805  0.671068
subsex:domneut:subneut:domage              0.00057    1.1   1    0.5148  0.473266
domsex:subsex:domneut:subneut:domage       0.00133    2.5   1    1.1951  0.274596
Residuals                                          1903.0 898
---
Signif. codes:  0 ‘***’ 0.001 ‘**’ 0.01 ‘*’ 0.05 ‘.’ 0.1 ‘ ’ 1
```

**Reduced Model**

```
> B <- lm(bcPower(diff,0.67) ~ order+domsex*subsex+subneut+domage+subsex:subneut, data=newdomdiff, na.action = na.omit)
> etasq(B, anova = TRUE, type=3)
Anova Table (Type III tests)

Response: bcPower(diff, 0.67)
               Partial eta^2 Sum Sq  Df   F value    Pr(>F)
(Intercept)          0.82978 9501.2   1 4499.2793 < 2.2e-16 ***
order                0.00907   17.8   1    8.4453 0.0037473 **
domsex               0.00081    1.6   1    0.7491 0.3869709
subsex               0.01545   30.6   1   14.4872 0.0001504 ***
subneut              0.00552   10.8   1    5.1186 0.0239021 *
domage               0.00709   13.9   1    6.5948 0.0103839 *
domsex:subsex        0.00483    9.5   1    4.4758 0.0346454 *
subsex:subneut       0.00532   10.4   1    4.9340 0.0265747 *
Residuals                    1949.1 923
---
Signif. codes:  0 ‘***’ 0.001 ‘**’ 0.01 ‘*’ 0.05 ‘.’ 0.1 ‘ ’ 1
> summary(B)

Call:
lm(formula = bcPower(diff, 0.67) ~ order + domsex * subsex + subneut +
    domage + subsex:subneut, data = newdomdiff, na.action = na.omit)

Residuals:
    Min      1Q  Median      3Q     Max
-4.1251 -0.9660 -0.0121  1.0361  3.8657

Coefficients:
                           Estimate Std. Error t value Pr(>|t|)
(Intercept)                  7.8793     0.1175  67.077  < 2e-16 ***
orderSecond                 -0.3978     0.1369  -2.906  0.00375 **
domsexmale                   0.1175     0.1358   0.866  0.38697
subsexmale                  -0.6133     0.1611  -3.806  0.00015 ***
subneutneutered             -0.3054     0.1350  -2.262  0.02390 *
domageyounger                0.3243     0.1263   2.568  0.01038 *
domsexmale:subsexmale        0.4127     0.1951   2.116  0.03465 *
subsexmale:subneutneutered   0.4340     0.1954   2.221  0.02657 *
---
Signif. codes:  0 ‘***’ 0.001 ‘**’ 0.01 ‘*’ 0.05 ‘.’ 0.1 ‘ ’ 1

Residual standard error: 1.453 on 923 degrees of freedom
Multiple R-squared:  0.03601,	Adjusted R-squared:  0.02869
F-statistic: 4.925 on 7 and 923 DF,  p-value: 1.77e-05

> Residuals <- residuals (B)
> qqnorm(Residuals)
> shapiro.test (Residuals)

	Shapiro-Wilk normality test

data:  Residuals
W = 0.99671, p-value = 0.05003
```

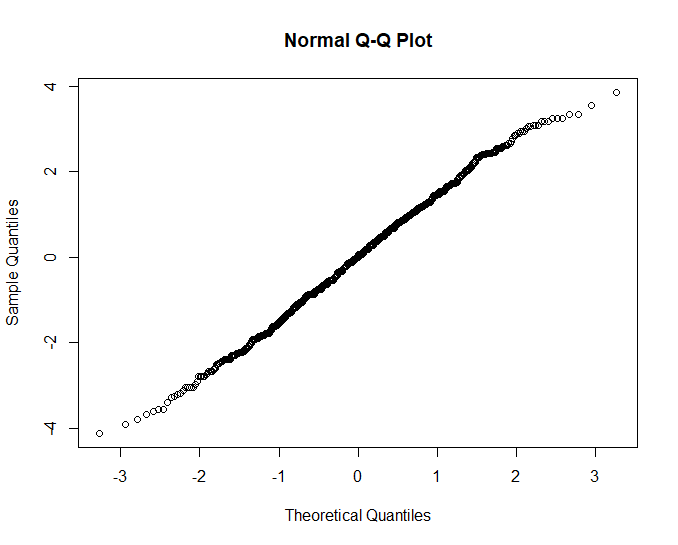
